# An *in silico* survey of *Clostridioides difficile* extrachromosomal elements

**DOI:** 10.1101/651539

**Authors:** Bastian Hornung, Ed J. Kuijper, Wiep Klaas Smits

## Abstract

The gram-positive enteropathogen *Clostridioides difficile* is the major cause of healthcare associated diarrhoea and is also an important cause of community-acquired infectious diarrhoea. Considering the burden of the disease, many studies have employed whole genome sequencing to identify factors that contribute to virulence and pathogenesis. Though extrachromosomal elements such as plasmids are important for these processes in other bacteria, the few characterized plasmids of *C. difficile* have no relevant functions assigned and no systematic identification of plasmids has been carried out to date. Here, we perform an *in silico* analysis of publicly available sequence data, to show that ∼13% of all *C. difficile* strains contain extrachromosomal elements, with 1-6 elements per strain. Our approach identifies known plasmids (e.g. pCD6, pCD630 and cloning plasmids) and 6 novel putative plasmid families. Our study shows that plasmids are abundant and may encode functions that are relevant for *C. difficile* physiology. The newly identified plasmids may also form the basis for the construction of novel cloning plasmids for *C. difficile* that are compatible with existing tools.

**Repositories:** The assembled circular type plasmids have been deposited at the European Nucleotide Archive (ENA) under accession numbers ERZ940801 and ERZ940803-ERZ940808.

## Introduction

*Clostridioides difficile* (*Clostridium difficile*) [1] is a gram-positive, endospore forming, anaerobic bacterium. It is an opportunistic pathogen in humans, and is the causative agent of most cases of antibiotic associated diarrhoea [2]. In recent years, the bacterium is also increasingly found in cases of infectious diarrhoea that cannot be linked to healthcare exposure [2]. *C. difficile* infections can be refractory to antimicrobial therapy and even when initial cure is observed, relapses are frequent [2]. Typing methods for *C. difficile* include (capillary) PCR ribotyping, multilocus sequence typing (MLST) and single nucleotide polymorphism (SNP) typing after whole genome sequencing [3]. Since the beginning of the 21^th^ century, an increase in *C. difficile* infections due to epidemic types such as PCR ribotype 027 and 078 has been noted [4, 5]. Although the molecular mechanisms underlying the epidemicity are poorly understood and remain under debate [6], robust toxin production and sporulation [7], altered surface properties [8], resistance to antimicrobials [5, 9] and an increased ability to metabolize certain sugars [10] have been implicated.

In other organisms, the contribution of plasmid-encoded functions to virulence and pathogenesis is well-documented [11-13]. By contrast, only a limited number of plasmids has been identified in *C. difficile* and all of these are cryptic, *i.e.* no traits have been associated with plasmid carriage. Commonly used cloning vectors for *C. difficile* make use of a replicon derived from the 6.8 kb plasmid pCD6 [14]. The reference strain 630 contains a single plasmid, pCD630, that is part of a larger family of 7.8-11.8 kb plasmids [15, 16]. And recently several large (>42 kb) plasmids were described [17]. Nevertheless, earlier work supports the notion that plasmids may be common in *C. difficile* [18, 19]. It is also possible, however, that other extrachromosomal elements were detected in these studies: both conjugative transposons and phages can exists as circular double stranded DNA intermediates [20-23].

Several approaches to identify plasmids from next generation sequencing data have been described, based on sequence homology, coverage, contig interactions or machine learning approaches [24]. None of these have been applied for a systematic investigation of extrachromosomal elements in *C. difficile*.

Here, we used a bioinformatic approach to identify extrachromosomal elements in publicly available *C. difficile* whole genome sequence data. Based on our analysis, we expect that most of these elements represent plasmids, suggesting a so far untapped potential for virulence determinants, or source for genetic engineering of diverse *C. difficile*.

## Methods

### Detection of putative extrachromosomal elements in public databases

To identify extrachromosomal elements in a high throughput manner, we implemented an approach comparable to `placnetW` [25, 26], a graph-based tool for reconstruction of plasmids from next generation sequence pair-end datasets (**Figure 1**). In short, 5403 public paired-end Illumina datasets were downloaded from the NCBI (for accession numbers see Supplemental Table 1; accessed on 07-09-2017). All samples were downloaded with `eutils` prefetch [27] and converted with `fastq-dump` from the SRA toolkit v2.8.2-1. The optimal kmer was predicted by `kmergenie` v1.6741 [28] on the interleaved fastq files and the assembly was performed with `Velvet` v1.2.10 [29]. Afterwards the assembly graph from the Velvet output was parsed into a graph with the python `networkX` library v1.11 [30]. To calculate size and coverage, all headers in the Velvet assembly were parsed into the network. The biggest component (based on size in bases) was considered to be the genome, and average coverage was estimated by averaging the coverage of all contigs over the amount of contigs. All other network components were considered to belong to the chromosome if their coverage did not exceed 1.5 times the coverage of the chromosome.

**Figure 1.**
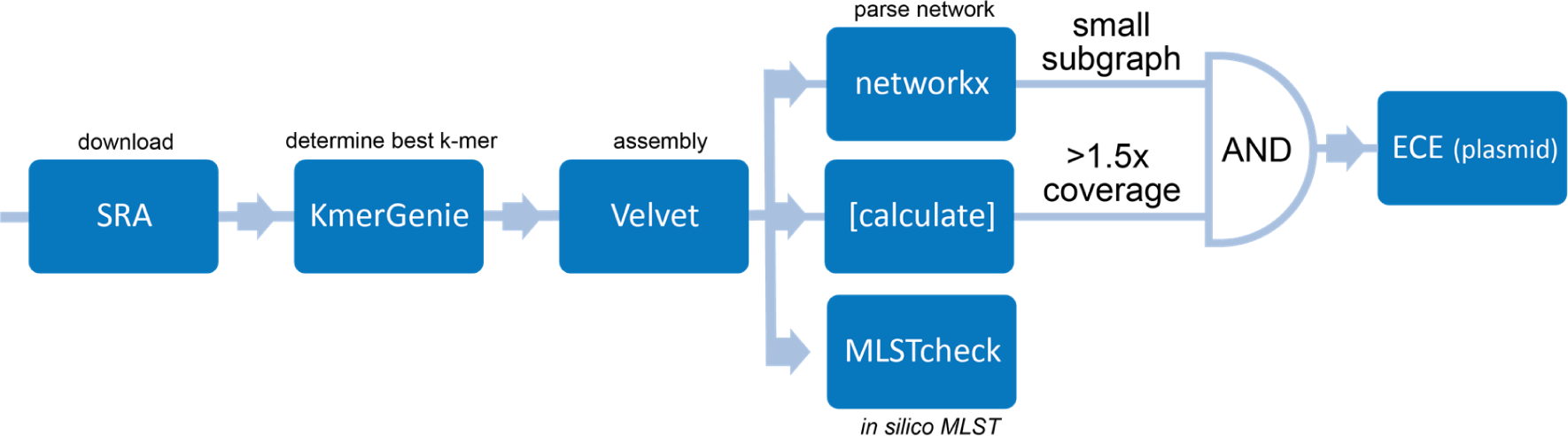
Approach. Schematic representation of the approach for the *in silico* identification of extrachromosomal elements in *C. difficile*.

To reduce the amount of false positive identifications, in a second step the coverage was adjusted for the number of base pairs instead of the number of contigs. Furthermore, a `Blast` search (v2.7.1) was performed against the chromosome of *Clostridium difficile* 630 [15], and all components with more than 50% genomic content were regarded as belonging to the chromosome. Additionally, a blast search was performed with all identified sequences against the NCBI plasmid database ([27]; download 11-09-2017).

To detect homology between the assembled sequences, the average nucleotide identity (ANI) was calculated between all plasmids with `pyANI` v0.2.7 [31]. The following known plasmids were included as well: pCD6 (AY350745.1), pCD630 (AM180356.1), pCD-WTSI1 (MG019959.1), the plasmid from *C. difficile* strain BI1 (FN668942.1), the big plasmids 1 and 2 of strain FDAARGOS_267 (NZ_CP020425.1 and NZ_CP020426.1), pAK1 and pAK2 (NZ_CP027015.1 and NZ_CP027016.1), pHSJD-312 (MG973074.1), pCd13_cfrC (MH229772.1), LIBA6289 (MF547664.1), pZJCDC-S82 (JYNK01000020.1), *Clostridioides difficile* strain CD161 plasmid unnamed1 and 2 (CP029155.1 and CP029156.1), *Clostridioides difficile* strain CDT4 plasmid unnamed1 (CP029153) and 25 described phages [20]. To determine the exact grouping, a further clustering analysis was performed. Per group, one representative was chosen, based on the results of the `pyANI` analysis (phi-X174, phiCD38-2, phiCD119, pCD-WTSI1, ERR1015479 plasmid #3 (ECE1), ERR251819 plasmid#2 (ECE2), ERR125924 plasmid #2 (ECE3), ERR347487 plasmid #2 (ECE4), ERR340291 plasmid #2 (ECE5), ERR251831 plasmid #1 (ECE6), as well as ERR125492 plasmid #1 (pCD-SMR), ERR247053 plasmid #1 (aminoglycoside resistance) and ERR125936 plasmid #1 (bacteriocin plasmid). Based on the ANI values, a hierarchical clustering with complete linkage (`Conda` v4.5.11 [32, 33], `Python` v2.7 [34], `SciPy` v1.0.0 [35]) was performed with these references and each plasmid. Each plasmid was assigned a type corresponding to its closest neighbour, unless the closest neighbour could not be exactly determined, or the branch rooted deeper in the tree.

Detailed comparative analysis of these sequences was performed with `Mauve` v2.3.1 [36] and `Blast` [37]. All Blast searches within this project were performed with the parameters –evalue 0.0001 and –culling_limit 1, unless otherwise mentioned.

Sequence typing of the assembled genomes was performed with `MLSTcheck` v2.1 [38].

Contamination checking of the assembled genomes was performed with `Mash` v2.0 [39] (with default parameters). Genomes were considered contaminated if any match with an e-value bigger than zero was present, which did not contain the words “*Clostridioides difficile*”, “*Clostridium difficile*”, “*Peptoclostridium difficile*”, “*Clostridium phage*”, “Enterobacteria phage phiX174” or “*Clostridium* sp. HMSC”. Entries containing the word “plasmid” were ignored.

### Plasmid annotation

Annotation was performed with another in-house pipeline. This pipeline consists out of gene calling with `prodigal` version v2.6.3 (with –meta option) [40], rRNA prediction with `RNAmmer` v1.2 [41], tRNA prediction with `Aragorn` 1.2.38 [42] and CRISPR annotation with the `CRISPR recognition tool` v1.2 [43]. Genes, which were predicted over an assembly gap and contained more than 50% N were discarded. Protein annotation was performed with `InterproScan` v5.26-65.0 [44], `PRIAM` vMarch 2015 [45] (together with legacy `Blast` v2.2.26 [37]) and `dbCAN` v5.0 [46]. Additional EC numbers were derived via the GO terms [47] derived from the InterproScan output. If an EC number could be assigned to a protein sequence, it was annotated with the canonical name of its EC number. Otherwise, all InterproScan domain names were searched for terms relating to functions involved in virus replication, sporulation, ribosomal proteins, CRISPR or any “subunit” containing names, and these names were used with priority for the naming. If this did not lead to any result, all domain names were searched for words ending in “ase”, indicating potential enzyme functions. Otherwise a random domain name was picked. All these annotation steps were done while disregarding generic or uninformative terms (e.g. containing “hypothetical”, “DUF”, “uncharacterized”). Domains containing these words were only considered after all other steps did not lead to any result. All programs were executed with standard parameters, unless specific parameters were mentioned.

### CRISPR analysis

CRISPR elements were predicted for all genomes with the `CRISPR recognition tool` v1.2 [43]. A blast database was built for all the different ECE groups based on the `pyANI` analysis. A blast search of all the CRISPR spacers against these databases was performed with the options -task blastn-short -outfmt “6 std qlen”, and afterwards all hits were filtered for having at least a match of 90% of the query length.

### Data accessibility

All reference plasmid assemblies have been uploaded to the European Nucleotide Archive under accession numbers ERZ940801, ERZ940803-ERZ940808

## Results

### Extrachromosomal elements are abundant in C. difficile

There is substantial evidence that plasmids are more abundant in *C. difficile* than expected on the basis of the published number of characterized plasmids from this organism [16-19]. We set out to determine the prevalence and identity of extrachromosomal elements (ECEs) in *C. difficile* in a high-throughput manner. To this end, we analysed public whole genome sequence data from the National Center for Biotechnology Information. 5403 samples (**Supplemental Table 1**) in the sequence read archive which were sequenced on Illumina machines in paired-end mode were processed using an in-house pipeline (**Figure 1**) based on `PLACNETw` [26]. Of these, 5336 genomes were successfully assembled. In total we identified 1066 putative extrachromosomal elements within 692 genomes, which corresponds to a prevalence of 13% (**Supplemental Table 2**). These data confirm that ECEs are abundant in *C. difficile*.

Most (451) of the genomes contained a single ECE, but the presence of two or three elements was also common (137 and 76 genomes respectively) (**Figure 2A**). The highest number of ECEs observed for a single genome was six. This indicates that at least some of the ECEs are compatible with others.

**Figure 2.**
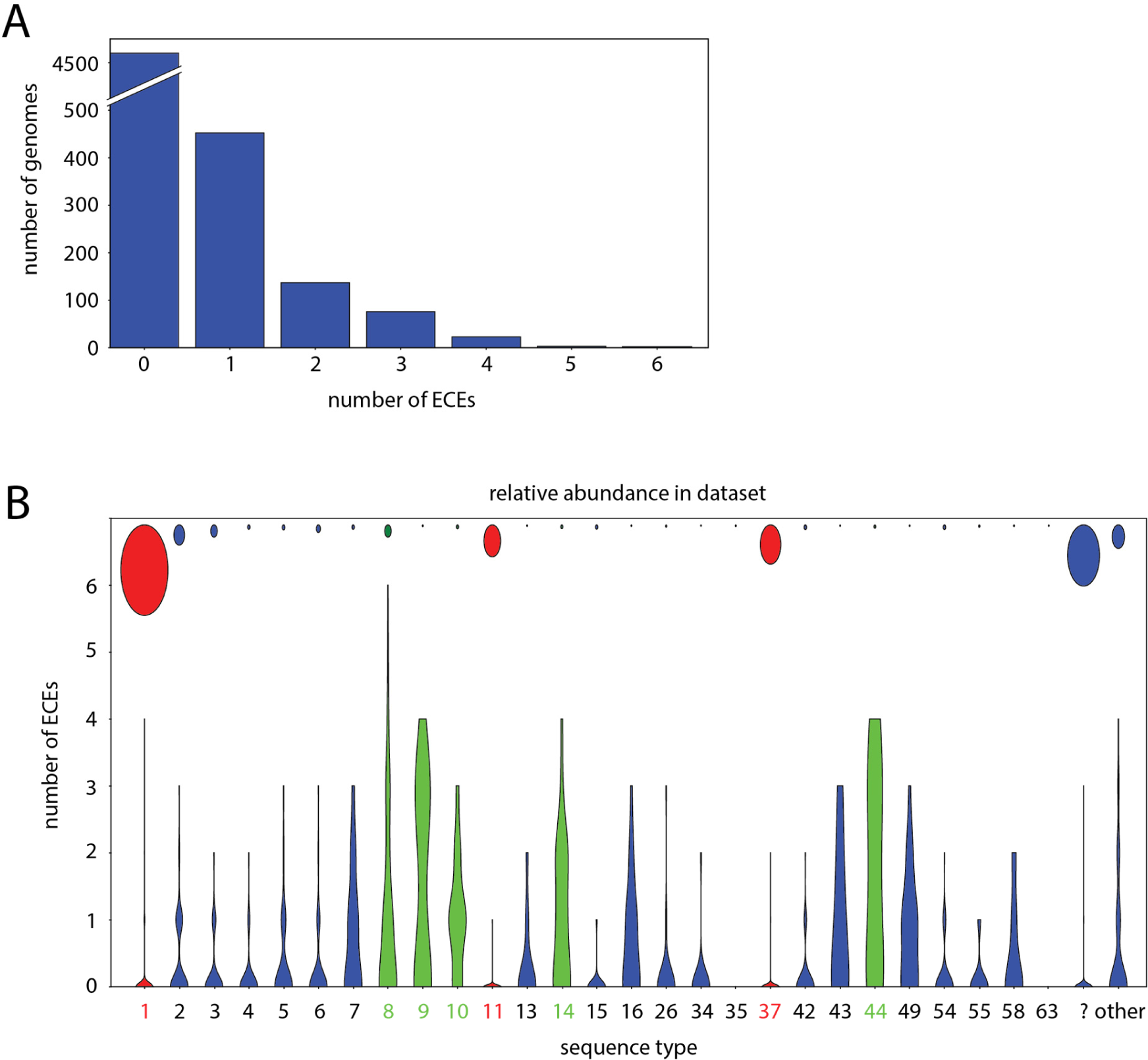
Extrachromosomal elements of *C. difficile*. **A**. Number of ECEs per strain. **B**. Distribution in the number of extrachromosomal elements amongst various sequence types of *C. difficile*. The violin plots indicate the distribution of plasmids for a particular sequence type (in the figure ST1/1* is indicated as “1”), whereas the circles indicate the relative amount of sequenced genomes of a particular sequence type. Only sequence types with 20 or more sequenced genomes are displayed separately. All other sequence types are summarized under “other”. Genomes to which no sequence type could be assigned are summarized under “?”. The sequence types belonging to the hypervirulent ribotypes RT27 (ST1) and RT078 (ST11), as well as the widely sequenced ST37 are marked in red. Sequence types with more than one plasmid per genome on average are marked in green.

### ECE content differs for different multilocus sequence types

Above, we observed variation in the number of ECEs per genome sequence. We wanted to establish whether certain types of *C. difficile* were more likely to contain ECEs than others. High quality whole genome sequence data allows the determination of sequence types (ST). We could successfully assign a ST to 52% (2770/5336) of the genome sequences using MLSTcheck (**Supplemental Table 1**). A substantial number of genomes could not be assigned a ST (n=914), or were designated as closely related to an identifiable sequence type (n=1652) using this tool. For the analyses hereafter, we will refer to the latter group using their closest sequence type (e.g. ST1 indicates ST1 only, whereas ST1/1* indicates both ST1 and and STs closely related to ST1).

Whole genome sequencing data is biased towards clinical isolates as these are most frequently investigated. This is reflected in the relative abundance in our dataset of the well-known epidemic PCR ribotypes RT027 and RT078, that were subject of extensive whole genome sequencing studies [5, 9]. RT027 and RT078 belong to ST1 (clade 2) and ST11 (clade 5), respectively. 1352 genomes were assigned to ST1/1*, and 477 to ST11/11*. The next largest group (586 genomes) corresponds to ST37/37* (that includes the toxin A negative PCR RT017, clade 4). Together, these three groups make up 45% of all the sequences analysed. Despite being so widely sequenced, ST1/1*, ST11/11* and ST37/37* contained only 62, 4 and 6 ECEs, respectively, of the 1066 elements identified here (**Figure 2B**).

Further analysis suggests that most of the ECEs in ST1/1*, ST11/11* and ST37/37* are in fact likely bacteriophages. These are in part derived from technical spike-in controls (phiX174) [48]: 21/62 of the ST1/1*, 0/4 of the ST11/11* and 1/6 of the ST37/37* ECEs correspond to this phage. *Clostridium* phage phiCD38-2 [49] was also common: 20/62 ST1/1*, 1/4 ST11/11* and 4/6 ST37/37* ECEs correspond to this phage. Finally, in ST11, *Clostridium* phage phiCD6356 [50] was identified once. Overall, 48/72 ECEs in these epidemic types are likely to be phage, further reducing the number of putative plasmids in these groups. Notably, the majority of ST1/11/37/* isolates does not contain any ECEs, suggesting a possible negative correlation between ECE carriage and epidemicity.

By contrast, we noticed that certain ST more frequently contain ECEs. For example, ST8/8* contained 189 ECEs in 183 analyzed genomes, with 1-6 per genome (**Figure 2B**). ST8 includes RT002, the 7^th^ most common PCR ribotype in Europe [51]. Other STs that appear to contain at least one ECE on average are ST9/9*, ST10/10*, ST14/14* and ST44/44* (**Figure 2B**). Notably, all these sequence types fall in clade 1 [52, 53]. The highest average ECE content was observed for ST9 with 39 elements in 15 samples (2.6/genome), followed by ST44 2.2/genome).

Taken together, our data suggest that certain *C. difficile* types may be more tolerant to plasmid carriage than others.

### pCD630- and pCD6-like plasmids are common

We wanted to confirm that our pipeline can identify *bona fide* plasmids. We therefore screened the identified ECEs against known *C. difficile* plasmid sequences, like pCD6, pCD630 and others (see Methods). We found several plasmids highly similar and sometimes identical to these known ones, derived from various STs. The class pCD-WTSI1/pCD630 like plasmids contained 378 ECEs (the majority more similar to pCD-WTSI1), and 189 plasmids were similar to pCD6 (these are contained in a larger family of 296 ECEs, see ECE6 below). We also identified various ECEs overlapping with the previously identified phages and megaplasmids (n=70) [17].

Interestingly, we inadvertently also identified a replicative cloning plasmid carrying chloramphenicol/thiamphenicol resistance gene in one of the whole genome sequences (ERR125924). Plasmids are generally introduced as shuttle vectors from the hosts *Escherichia coli* [14] or *Bacillus subtilis* [54]. We screened all ECEs for regions associated required for transfer from the conjugation donor (*traJ*/*oriT* or the Tn*916 oriT)* and replication in gram-negatives (pBR322/ColE1 origins), but did not find any further cloning vector contaminations.

Overall, our data confirms that our analysis does in fact detect *bona fide* plasmids, as well as certain phages. Plasmids with significant homology to the characterized plasmids pCD630 [15, 16] and pCD6 [14] are the most common (**Supplemental Table 2**).

### Identification of 6 novel families of ECEs

Many of the ECEs identified do not have homology to already described phages and plasmids discussed above (n=222), or share only limited homology (n=107). We reasoned that those ECEs that are part of a family are more likely to represent legitimate *C. difficile* plasmids and therefore clustered the ECEs by sequence similarity. This resulted in the identification of 6 putative families of ECEs (n=478), and a group of singletons (n=40) (**Supplemental Table 2**). Some details of the different families are discussed below.

#### ECE1

The 6.1kb type plasmid pCD-ECE1 (ERZ940803) is derived from ERR1015479 (**Figure 4A**). Plasmids in this family range in size between 6071 and 7284bp and appeared in 21 samples. The distribution of STs showed some clustering, with seven samples belonging to ST436, and coming from the bioproject PRJEB5486, where eight samples were sequenced. The eighth sample of this bioproject also contained this plasmid, but it was assigned to ST9 (and potentially contaminated with a *Lactobacillus*). Another 10 more samples belonged to ST9, 2 ST10 and 1 ST75*. 9 of these 21 sequences were predicted to be circular with eight of these having a length of 6071bp and a minimum identity over the full length of 99.9%.

**Figure 3.**
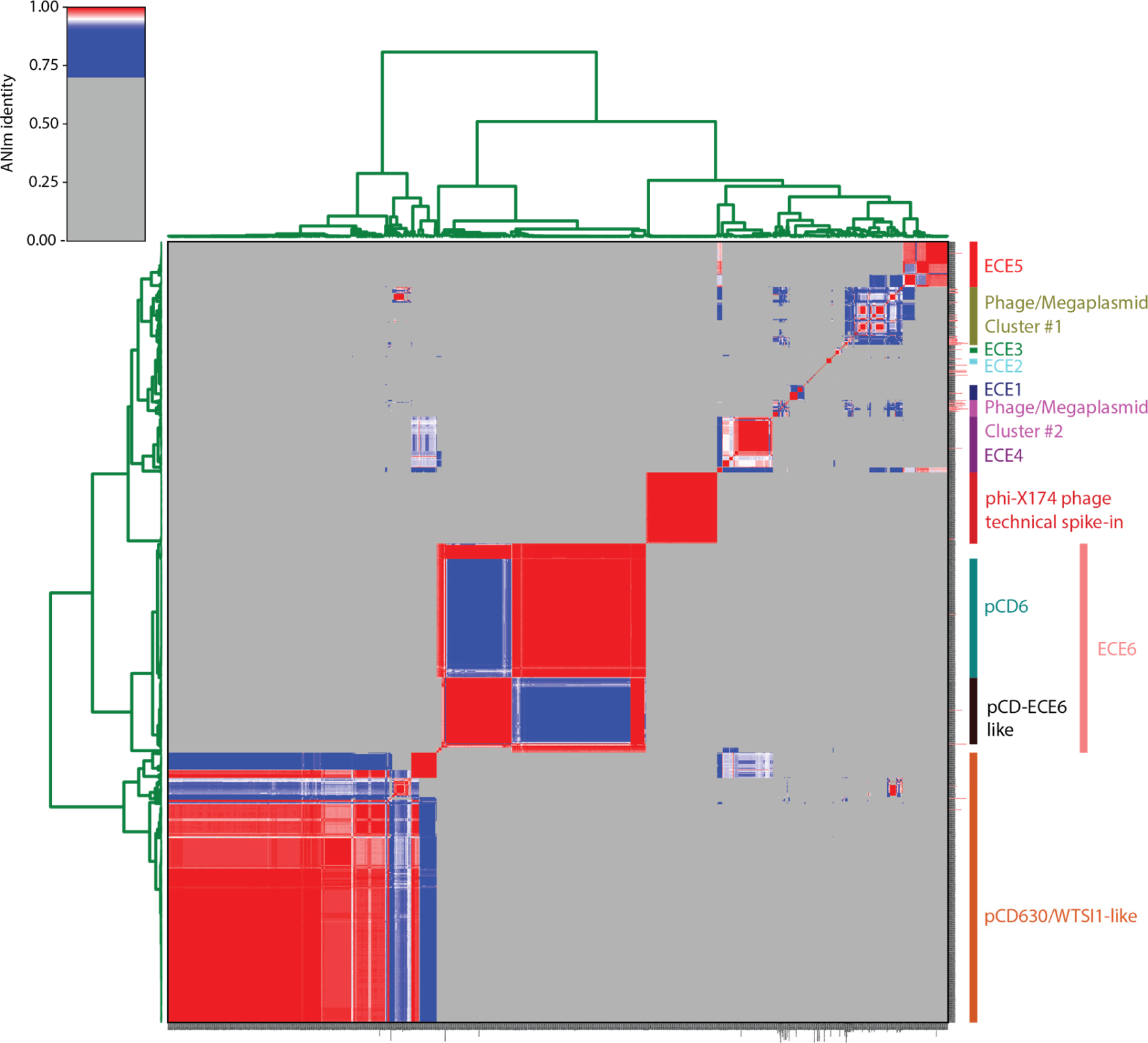
Identification of ECE families. Hierarchical clustering based on average nucleotide identity was performed. Red to blue colors indicate similarity. Gray values indicate that no significant similarity was found. Names of the clusters are given on the right. A high resolution version of this figure (with readable plasmid names) is provided as Supplemental

**Figure 4.**
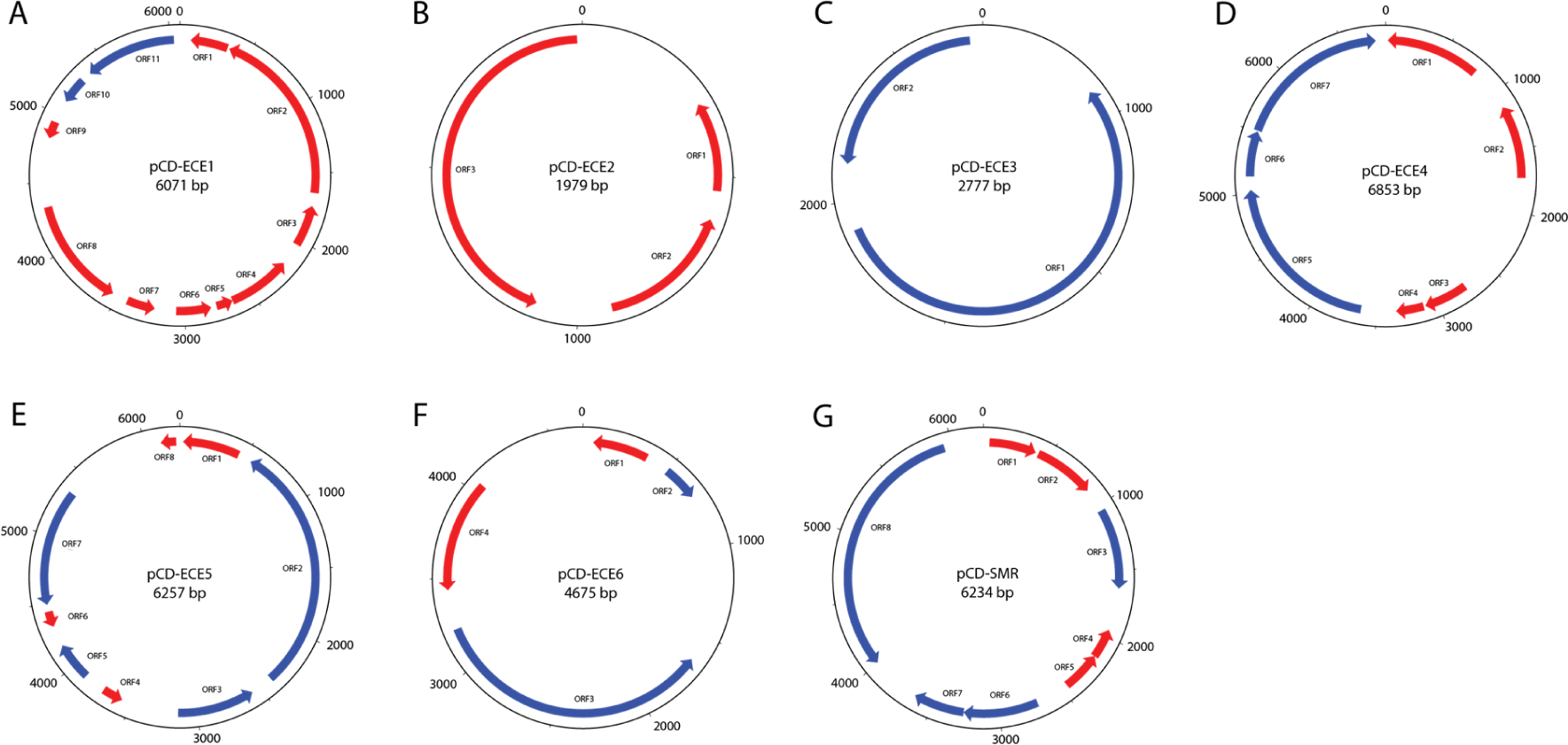
Maps of circular extrachromosomal elements. Schematic representation of representative members of the 6 putative plasmid families pCD-ECE1 (A), pCD-ECE2 (B), pCD-ECE3 (C), pCD-ECE4 (D), pCD-ECE5 (E) and pCD-ECE6 (F), and the singleton putative multidrug resistance plasmid, pCD-SMR (G). Ticks indicate 500bp intervals, with every 1000bp labelled. Red open reading frames are hypothetical (no results in automated annotation pipeline, see Materials and Methods). Blue open reading frames indicate genes for which a putative function could be assigned in automated annotation. **A**. ORF10: Arc-type ribbon-helix-helix protein. ORF11: Zonular occludens toxin. **B**. No functions annotated. **C**. ORF1: plasmid replication protein, ORF2: plasmid recombination enzyme. **D**. ORF5: Plasmid recombination enzyme, ORF6: penicillinase repressor, ORF7: BlaR1 peptidase M56. **E**. ORF2: Type III restriction enzyme (res subunit), ORF3: phage integrase family protein, ORF5: HNH endonuclease 5, ORF7: DNA-directed DNA polymerase. **F**. ORF2: Lambda repressor-like, DNA-binding domain superfamily protein, ORF3: alpha-helical domain, primase C-terminal like. **G**. ORF3: Winged-helix-like DNA-binding domain superfamily protein, ORF6: Bacterial regulatory protein, tetR family, ORF7: Small multidrug resistance protein, ORF8: MobA/MobL protein.

#### ECE2

The 2.0kb type plasmid pCD-ECE2 (ERZ940804) is derived from ERR251819 (**Figure 4B**). Plasmids in this family appeared in 7 samples belonging to 5 STs from 5 different bioprojects. Six of these were assembled into a single contig of 1979bp, and were due to overlaps confirmed to be circular. Nearly all plasmids showed 100% identity, with only a single SNP in one of the plasmids. The 7^th^ plasmid was fragmented into multiple contigs, which overlapped with each other, with a cumulative non-redundant sum of 1979bp. Also this plasmid showed a minimum 99% identity to the other plasmids. Functional annotation only showed 3 hypothetical proteins on each plasmid. No significant homology to any sequence in the NCBI database could be found.

#### ECE3

The 2.8kb type plasmid pCD-ECE3 (ERZ940805) is derived from ERR125924 (**Figure 4C**). Plasmids in this family were found in 10 samples, of which five belonged to ST1/1*. Four of these were predicted to be circular and their size varied between 2.8 and 6.8kb. 2 plasmids, derived from different STs within the same bioproject, were circular and 100% identical with a size of 3361bp. They showed a total homology of >2000bp with an identity of 94-95% to the reference plasmid. Homology was less strong to the 4^th^ circular plasmid, which had only an identity of 86-88% over a length of 1414/460bp. Functional annotation only identified plasmid replication and recombination proteins, in addition to hypothetical proteins, and in one case a DNA polymerase.

#### ECE4

The 6.9kb type plasmid pCD-ECE4 (ERZ940806) is derived from ERR347487 (**Figure 4D**). The 79 plasmids in this family were found in samples from more than 10 different STs from various bioprojects. The size ranged from 5kb to 22kb. These plasmids include 4 identical circular plasmids with a size of 15kb, belonging to ST9/9* from 2 different bioprojects, and 21 circular plasmids with a size of 6853bp, belonging to >5 bioprojects and STs, and a few other identical plasmids with varying sizes. The diversity of sizes in the ECE4 group, based on our `pyANI` analysis, is striking. It is possible that more detailed characterization will reveal the existence of subfamilies or multiple families that have been grouped based on our method.

#### ECE5

The 6.3kb type plasmid pCD-ECE5 (ERZ940807) is derived from ERR340291 (**Figure 4E**). This family contains 65 plasmids from various STs and bioprojects. The size ranges from 5044 bp up to 12kb. This class includes 15 predicted circular plasmids with a size of 6257bp (present only in STs 6,7 and 8/8*), 8 circular plasmids of 6651 bp (belonging to ST 9, 12 and 436) and 4 circular plasmids of 5819 bp.

#### ECE6

The 4.7kb type plasmid pCD-ECE6 (ERZ940808) is derived from ERR251831 (**Figure 4F**). This family includes the well-characterized pCD6 and comprises 296 plasmids. We do not group plasmids with high similarity to pCD6 separately, as it cannot be clearly distinguished from the other plasmids in the `pyANI` analysis (**Figure 3**). It can be divided into 3 subfamilies: a small group (including ERR340281.plasmid_1, n=20) seemingly consisting of parts, a group that includes pCD6, and a further not yet characterized type of plasmid (pCD-ECE6). On the basis of the `pyANI` analysis, these subfamilies clearly group together (**Figure 3**). The size ranges from a single plasmid of 3335bp (16 contigs), through a group of circular plasmids with a size of 4675bp (n=45, belonging to various (>5) bioprojects and STs), up to a size of 18kb. Most of these sequences could not be confirmed to be circular, but various instances with varying sizes appeared in more than one sample with >95% identity. The largest that could be circularized had a size of 8297bp.

These 6 classes represent 478 of the 1066 identified ECEs. As indicated above already, the known class of pCD-WTSI1/pCD6 like plasmids contains 378 ECEs (majority more similar to pCD-WTSI1). The technical spike-in controls (phiX174) contain another 100 sequences. In total 70 sequences were assigned to probable *Clostridium* phages and megaplasmids (pCDBI1, pAK1, pAK2, etc).

40 ECEs do not fall into one of these families (singletons); these may represent legitimate low-prevalence plasmids in our sample population, or may reflect incidental contaminations. 2 known plasmids could not be classified into one of the groups either, pZJCDC-S82 (JYNK01000020.1) and pCd13_cfrC (MH229772.1), indicating that low-prevalence plasmids do exist in the population.

### Function of new ECE families

We annotated all these newly identified ECEs to be able to analyze their function. Many of the genes were annotated as coding for hypothetical proteins. Of the genes which had a function assigned, mostly functions related to DNA processing and plasmid replication (e.g. RNA polymerase, winged helix-like DNA-binding domain superfamily, helicase, resolvase, zinc fingers) or phage elements (e.g. integrase, phage tail, phage capsid) were present. Some of these functions and their relationships to different plasmid classes are detailed below.

### Novel ECE families may encode virulence factors

pCD-ECE1 belongs to a family of ECEs that is characterized by the presence of an Arc-type ribbon-helix-helix domain (IPR013321; ORF10) and an open reading frame with strong homology to the gene encoding the zonula occludens toxin (Zot, IPR008900; ORF11) (**Figure 4A**). Zot is an enterotoxin from bacteriophages infecting Proteobacteria, including enteropathogens such as *Vibrio* and *Campylobacter* spp, that increases intestinal permeability by affecting tight junctions and a relation with IBD has been suggested [55, 56]. Indeed, one of the ECEs from this family contained a gene coding for a phage capsid protein (IPR008020), which was not seen on the other plasmids. Notably, this plasmid showed only partial homology to the other ECE1 plasmids, with identities ranging between 84-93% over 2182 bp. Other obvious phage elements could not be identified in these ECEs. To none of the other genes any function could be assigned.

The contribution of this putative toxin to *C. difficile* infection, when carried on a plasmid, is currently unknown.

### Novel ECEs may encode antibiotic resistance determinants

A singleton ECE (not falling into one of the ECE families described above) was identified as carrying a putative antibiotic resistance determinant. This circular 6.2kb plasmid, dubbed pCD-SMR (derived from ERR125942; accession ERZ940801, ST1; **Figure 4G**) contains a gene encoding a TetR-family regulator (IPR001647; ORF6) adjacent to an small multidrug resistance protein (IPR000390; ORF7). It also contains a MobA/MobL mobilization protein (IPR005053; ORF8), and a winged helix-like DNA-binding domain superfamily protein (IPR036388; ORF3). A region of ∼1kb, containing part of the regulator and the drug transporter, may be derived from a *Listeria* transposon or plasmid pK5 from an uncultured organism (KJ792090.1).

Another singleton ECE (derived from sample ERR247053, ST48; **Supplemental Table 2**) was identified to contain a aminoglycoside-2’’-adenylyltransferase (IPR019646), that can provide resistance to the most common aminoglycosides (except streptomycin) and has been found in plasmids before [57]. The ECE was assembled in 9 contigs with a cumulative size of 5384 bp, only containing one more annotated open reading frame, encoding a mobilization protein. The contigs showed a high identity (>93%) to various genomic and plasmid regions of *Campylobacter jejuni*.

The majority the plasmids belonging to the ECE4 and ECE6 families (pCD6-like) carry the genes BlaR1 (IPR008756) and BlaI (IPR005650) (62/79 ECE4 both genes, 197/296 BlaR1 in ECE6 and 194/296 BlaI in ECE6). While these do not confer resistance to any antibiotics, they regulate the induction of beta-lactamases in various species [58, 59], including *C. difficile* [60]. The genes found on ECE4 and ECE6, however, show only limited similarities (20-30%) to the chromosomal regulators identified in that study [60]. Therefore the significance of the carriage of these genes for resistance to beta-lactams is currently unknown.

### Novel ECEs might potentially contain bacteriocins

One singleton ECE (derived from sample ERR125936, ST unknown; **Supplemental Table 2)** contained many functional domains unique to our ECE dataset. This ECE contained 3 contigs with a total size of 18812bp, and 20 ORFs. Despite very limited homology at the nucleotide level, most protein sequences showed homology to protein sequences derived from a single contig of approximately 19kb from an unknown *Muribaculaceae* species (RIAY01000031.1). The domains of interest include a peptidase of the C39 family, a TonB-dependent receptor, an ubiquitin-activating enzyme, an acyl-CoA N-acyltransferase, a protein with a generic prokaryotic membrane lipoporotein attachment site and a radical SAM enzyme (potentially split into two CDS over two contigs). We suggest this cluster to be involved in bacteriocin synthesis for three reasons. First, the Interpro descriptions of the peptidase (IPR005074) and the radical SAM enzyme (IPR023885) indicate a possible involvement in bacteriocin processing and biosynthesis of ribosomally synthesized and post-translationally modified peptides (RiPP) [61], respectively. Second, an analysis with `Bagel4` indicated that the ubiquitin-activating enzyme had homology to a putative bacteriocin biosynthesis protein from *Streptococcus thermophilus* (Q5LXQ2), and labelled the contig as potentially containing a sactipeptide [62]. Finally, TonB-dependent receptors, lipoproteins, radical SAM enzymes are common components of bacteriocin biosynthesis clusters and acylation is often observed during bacteriocin biosynthesis [63, 64]. While no specific bacteriocin gene could be pinpointed, the presence of most of the unusual genes on this ECE can be explained the presence of such a RiPP gene cluster.

### CRISPR spacers mainly target phage, not novel ECE families

Since some of the ECEs were identified as phages, we wondered if CRISPR-based resistance mechanisms against ECEs exist, and if these do or do not correlate with sequence type. We predicted 371008 CRISPR spacers in all genomes (n=3994; in 1340 genomes no CRISPR spacers could be predicted), making a total of 31968 unique CRISPR spacers. The highly sequenced sequence types ST1/1*, ST11/11* and ST37/37*, which do not seem to carry many ECEs (**Figure 2B**), showed on average a medium (94), high (134) and low (35) carriage of CRISPR spacers. The MLST types, which seem on average to carry ECEs (ST8/8*, ST9/9*, ST10/10*, ST14/14*, ST44/44*), showed medium and high numbers of CRISPRS (84, 83,105, 127 and 127, respectively). Thus, we did not see a clear relationship between ECE carriage and the number of CRISPR spacers.

We furthermore tried to match the CRISPR spacers against the various ECEs. We found only 5 CRISPR spacers (of 31968 unique CRISPR spacers), which matched against the technical spike-in phi-X174 phage, indicating that most of the prediction did not cause false positive artifacts.

In 110 cases CRISPR spacers matched against ECEs isolated from their own genome. 57 of these 110 were directed against ECEs of the class phiCD38-2, 32 against pCD630/pCD-WTSI1 family and 10 against ECE1. Thus, only a minor fraction of CRISPR spacers targets endogenous extrachromosomal elements.

Most CRISPR spacers appear to target the 2 classes of phage/megaplasmids (7387 and 3496 CRISPR spacers, respectively), whereas much less targeted the 6 novel ECE families (934, 29, 0, 112, 13, 148, respectively) or the pCD630/pCD-WTSI1 family (n=978). Moreover, the phage/megaplasmid-targeting CRISPR spacers seemed to be more broadly distributed (the most widespread CRISPR spacers for both clusters predicted in 651 and 711 genomes) than the CRISRP spacers targeting the other ECE classes (most widespread: 575, 64, 498, 22, 524, 632 for ECE1, 2, 4, 5, 6 and pCD630/pCD-WTSI1, respectively). Together these data suggest that CRISPR-mediated defense likely does not play a major role in resistance against the ECE families identified here.

## Discussion

Here, we present the first comprehensive an *in silico* survey of ECEs in *C. difficile*. Our major findings are that ECEs are abundant (∼13% of all genomes analysed), strains can simultaneously carry 2-6 ECEs from different families, and that there appear to be at least 6 families of ECEs that have not been characterized yet. The classification into families should direct and facilitate future functional analysis of the ECEs. For instance, epidemiological analyses and cloning of the putative toxin in a *C. difficile* shuttle vector and comparison with a non-toxin carrying will elucidate whether *zot* can contribute to disease severity in humans and animal models.

Our study has several limitations. First, due to the use of publicly available sequencing data without access to the strains from which these data were generated, we are unable to confirm that the identified ECEs are in fact plasmids. However, our work allows the identification of conserved regions in each ECE family, which can be further developed into plasmid-specific PCRs to screen available collections of strains. Plasmid-content of PCR-positive strains can then be verified using established methods [16, 17]. Next, due to a bias in the available genome sequences towards clinical isolates of specific PCR ribotypes, our analysis may fail to capture the full breadth of plasmids that are present in *C. difficile*. However, one might argue that for clinical relevance, the current collection should provide ample information. More targeted sequencing strategies, aimed at a greater diversity of *C. difficile* strains from underrepresented sources and geographic locations, may lead to the identification of more plasmids. Furthermore, the produced assemblies did not undergo any quality control, and are partially of substandard quality. The low quality assemblies fall mostly in the category to which no ST could be assigned, and therefore does not affect the analysis of genomes with a ST assigned. Since our identified plasmids are mainly of small size (<20 kb), and the majority belongs to one of the previously or newly identified classes, we do not expect that many of these are false positives. We rather expect that due to the poor quality some of the low copy megaplasmids might have been incorrectly lumped into the genome, and that the actual diversity in ECEs might be even greater than what we discovered. Finally, we note that the *in silico* methodology employed here clearly also identifies phages, that can exist as extrachromosomal dsDNA [20, 21, 65] (see for instance the phage/megaplasmid cluster in **Figure 3**). Nevertheless, plasmids may encode phage proteins (incomplete phages) as the result of recombination between plasmids and phage [16, 17, 66]. Though we show that our approach can accurately identify *bona fide* plasmids, establishing whether the identified ECEs are phage, plasmid or an intermediate may need additional experimentation.

An intriguing finding is that *C. difficile* can harbour multiple ECEs simultaneously in the absence of selection. Indeed, the finding that 630Δ*erm* can harbour both pCD6-based replicons and the native pCD630 plasmid [16] supports this notion. Our work identified a ST8 strain that contained 6 ECEs; these include a pCD6-like plasmid, a pCD-WTSI1-like plasmid (from the pCD630 family) and pCD-BI1-like plasmid. This suggests that there is no plasmid incompatibility, or that the co-occurring plasmids do not belong to the same compatibility group. So far, no experiments have been described that employ two plasmids with different replicons to simultaneously introduce or express two different constructs in *C. difficile.* Moreover, to our knowledge, genetic manipulation of *C. difficile* has been limited to a small set of PCR ribotypes to date (RT012, RT027 and RT078) and the efficiency of the different replicons has been found to differ between these ribotypes [67]. The ECE pool described here therefore represents an interesting possibility to adapt or expand existing tools for use in a broader range of *C. difficile* types.

For conjugative elements (CTn’s) transfer between *C. difficile* is well established [22] and transfer of the pathogenicity locus via a so-far uncharacterized mechanism has also been described [68]. For the ECEs identified in this study, it is unknown if they are transferable between strains. Though conjugative plasmids are generally larger than mobilizable plasmids [69], large plasmids in *C. difficile* are likely to be non-conjugative due to the absence of conjugation or mobilization functions [17]. In general, the ECEs identified in the present study are smaller (967 of 1066 ECEs are <20 kb) and certain ECEs do seem to encode proteins relevant for mobilization or conjugation (such as ORF8 of pCD-SMR, **Figure 4G**). It remains to be established if these functions allow inter-or intraspecies transfer from *C. difficile* and if so, what other requirements for transfer may exist.

Overall, this work provides a starting point for investigations into the role of plasmids in *C. difficile* physiology and opens up the possibility of generating novel cloning vectors that may be particularly suitable for the manipulation of one or more *C. difficile* types.

## Author statements

### Author contributions

Conceptualization: WKS. Methodology: BVHH. Software: BVHH. Validation: BVHH, WKS. Formal analysis: BVHH, WKS. Investigation: BVHH, WKS. Resources: WKS, EJK. Data curation: BVHH. Writing – Original Draft Preparation: BVHH, WKS. Writing – Review and editing: BVHH, EJK, WKS. Visualization: BVHH. Supervision: EJK, WKS. Project administration: WKS. Funding: EJK, WKS.

## Acknowledgements

The authors would like to thank Peter van ’t Hof and Leon Mei (Sequencing Analysis Support Core, LUMC) for their help with using the `biopet` pipelines and for helpful discussions. We thank Jeroen Corver for helpful discussions and critical reading of the manuscript.

## Data

ERZ940801 and ERZ940803-ERZ940808.

## Supplemental Information

**Supplemental Figure 1**. High resolution version of Figure 3.

**Supplemental Table 1**. Overview of Sequence Read Archive entries analysed for this study.

**Supplemental Table 2**. Overview of extrachromosomal elements of *C. difficile*.

## References

1. Oren A, Rupnik M. Clostridium difficile and Clostridioides difficile: Two validly published and correct names. Anaerobe. 2018;52:125–6. doi: 10.1016/j.anaerobe.2018.07.005. PubMed PMID: 30031828.

2. Smits WK, Lyras D, Lacy DB, Wilcox MH, Kuijper EJ. Clostridium difficile infection. Nat Rev Dis Primers. 2016;2:16020. doi: 10.1038/nrdp.2016.20. PubMed PMID: 27158839; PubMed Central PMCID: PMCPMC5453186.

3. Knetsch CW, Lawley TD, Hensgens MP, Corver J, Wilcox MW, Kuijper EJ. Current application and future perspectives of molecular typing methods to study Clostridium difficile infections. Euro Surveill. 2013;18(4):20381. PubMed PMID: 23369393.

4. Goorhuis A, Bakker D, Corver J, Debast SB, Harmanus C, Notermans DW, et al. Emergence of Clostridium difficile infection due to a new hypervirulent strain, polymerase chain reaction ribotype 078. Clin Infect Dis. 2008;47(9):1162–70. doi: 10.1086/592257. PubMed PMID: 18808358.

5. He M, Miyajima F, Roberts P, Ellison L, Pickard DJ, Martin MJ, et al. Emergence and global spread of epidemic healthcare-associated Clostridium difficile. Nature genetics. 2013;45(1):109–13. doi: 10.1038/ng.2478. PubMed PMID: 23222960; PubMed Central PMCID: PMCPMC3605770.

6. Smits WK. Hype or hypervirulence: a reflection on problematic C. difficile strains. Virulence. 2013;4(7):592–6. doi: 10.4161/viru.26297. PubMed PMID: 24060961; PubMed Central PMCID: PMCPMC3906292.

7. Merrigan M, Venugopal A, Mallozzi M, Roxas B, Viswanathan VK, Johnson S, et al. Human hypervirulent Clostridium difficile strains exhibit increased sporulation as well as robust toxin production. J Bacteriol. 2010;192(19):4904–11. doi: 10.1128/JB.00445-10. PubMed PMID: 20675495; PubMed Central PMCID: PMCPMC2944552.

8. Vedantam G, Clark A, Chu M, McQuade R, Mallozzi M, Viswanathan VK. Clostridium difficile infection: toxins and non-toxin virulence factors, and their contributions to disease establishment and host response. Gut Microbes. 2012;3(2):121–34. doi: 10.4161/gmic.19399. PubMed PMID: 22555464; PubMed Central PMCID: PMCPMC3370945.

9. Knetsch CW, Kumar N, Forster SC, Connor TR, Browne HP, Harmanus C, et al. Zoonotic Transfer of Clostridium difficile Harboring Antimicrobial Resistance between Farm Animals and Humans. J Clin Microbiol. 2018;56(3). doi: 10.1128/JCM.01384-17. PubMed PMID: 29237792; PubMed Central PMCID: PMCPMC5824051.

10. Collins J, Robinson C, Danhof H, Knetsch CW, van Leeuwen HC, Lawley TD, et al. Dietary trehalose enhances virulence of epidemic Clostridium difficile. Nature. 2018;553(7688):291–4. doi: 10.1038/nature25178. PubMed PMID: 29310122; PubMed Central PMCID: PMCPMC5984069.

11. Adams V, Li J, Wisniewski JA, Uzal FA, Moore RJ, McClane BA, et al. Virulence Plasmids of Spore-Forming Bacteria. Microbiol Spectr. 2014;2(6). doi: 10.1128/microbiolspec.PLAS-0024-2014. PubMed PMID: 26104459.

12. Johnson TJ, Nolan LK. Pathogenomics of the virulence plasmids of Escherichia coli. Microbiol Mol Biol Rev. 2009;73(4):750–74. doi: 10.1128/MMBR.00015-09. PubMed PMID: 19946140; PubMed Central PMCID: PMCPMC2786578.

13. Pilla G, Tang CM. Going around in circles: virulence plasmids in enteric pathogens. Nat Rev Microbiol. 2018;16(8):484–95. doi: 10.1038/s41579-018-0031-2. PubMed PMID: 29855597.

14. Purdy D, O’Keeffe TA, Elmore M, Herbert M, McLeod A, Bokori-Brown M, et al. Conjugative transfer of clostridial shuttle vectors from Escherichia coli to Clostridium difficile through circumvention of the restriction barrier. Mol Microbiol. 2002;46(2):439–52. PubMed PMID: 12406220.

15. Sebaihia M, Wren BW, Mullany P, Fairweather NF, Minton N, Stabler R, et al. The multidrug-resistant human pathogen Clostridium difficile has a highly mobile, mosaic genome. Nat Genet. 2006;38(7):779–86. Epub 2006/06/29. doi: 10.1038/ng1830. PubMed PMID: 16804543.

16. Smits WK, Weese JS, Roberts AP, Harmanus C, Hornung B. A helicase-containing module defines a family of pCD630-like plasmids in Clostridium difficile. Anaerobe. 2018;49:78–84. doi: 10.1016/j.anaerobe.2017.12.005. PubMed PMID: 29246842.

17. Amy J, Bulach D, Knight D, Riley T, Johanesen P, Lyras D. Identification of large cryptic plasmids in Clostridioides (Clostridium) difficile. Plasmid. 2018;96-97:25–38. doi: 10.1016/j.plasmid.2018.04.001. PubMed PMID: 29702124.

18. Muldrow LL, Archibold ER, Nunez-Montiel OL, Sheehy RJ. Survey of the extrachromosomal gene pool of Clostridium difficile. J Clin Microbiol. 1982;16(4):637–40. PubMed PMID: 7153313; PubMed Central PMCID: PMCPMC272436.

19. Clabots C, Lee S, Gerding D, Mulligan M, Kwok R, Schaberg D, et al. Clostridium difficile plasmid isolation as an epidemiologic tool. Eur J Clin Microbiol Infect Dis. 1988;7(2):312–5. PubMed PMID: 3134239.

20. Fortier LC. Bacteriophages Contribute to Shaping Clostridioides (Clostridium) difficile Species. Front Microbiol. 2018;9:2033. doi: 10.3389/fmicb.2018.02033. PubMed PMID: 30233520; PubMed Central PMCID: PMCPMC6127314.

21. Sekulovic O, Fortier LC. Characterization of Functional Prophages in Clostridium difficile. Methods Mol Biol. 2016;1476:143–65. doi: 10.1007/978-1-4939-6361-4_11. PubMed PMID: 27507339.

22. Brouwer MS, Warburton PJ, Roberts AP, Mullany P, Allan E. Genetic organisation, mobility and predicted functions of genes on integrated, mobile genetic elements in sequenced strains of Clostridium difficile. PloS one. 2011;6(8):e23014. doi: 10.1371/journal.pone.0023014. PubMed PMID: 21876735; PubMed Central PMCID: PMCPMC3158075.

23. Mullany P, Allan E, Roberts AP. Mobile genetic elements in Clostridium difficile and their role in genome function. Res Microbiol. 2015;166(4):361–7. doi: 10.1016/j.resmic.2014.12.005. PubMed PMID: 25576774; PubMed Central PMCID: PMCPMC4430133.

24. Arredondo-Alonso S, Willems RJ, van Schaik W, Schurch AC. On the (im)possibility of reconstructing plasmids from whole-genome short-read sequencing data. Microb Genom. 2017;3(10):e000128. Epub 2017/11/28. doi: 10.1099/mgen.0.000128. PubMed PMID: 29177087; PubMed Central PMCID: PMCPMC5695206.

25. Lanza VF, de Toro M, Garcillán-Barcia MP, Mora A, Blanco J, Coque TM, et al. Plasmid Flux in Escherichia coli ST131 Sublineages, Analyzed by Plasmid Constellation Network (PLACNET), a New Method for Plasmid Reconstruction from Whole Genome Sequences. PLOS Genetics. 2014;10(12):e1004766. doi: 10.1371/journal.pgen.1004766.

26. Vielva L, de Toro M, Lanza VF, de la Cruz F. PLACNETw: a web-based tool for plasmid reconstruction from bacterial genomes. Bioinformatics. 2017. Epub 2017/10/17. doi: 10.1093/bioinformatics/btx462. PubMed PMID: 29036591.

27. Coordinators NR. Database resources of the National Center for Biotechnology Information. Nucleic Acids Res. 2013;41(Database issue):D8–D20. Epub 2012/11/30. doi: 10.1093/nar/gks1189. PubMed PMID: 23193264; PubMed Central PMCID: PMCPMC3531099.

28. Chikhi R, Medvedev P. Informed and automated k-mer size selection for genome assembly. Bioinformatics. 2014;30(1):31–7. Epub 2013/06/05. doi: 10.1093/bioinformatics/btt310. PubMed PMID: 23732276.

29. Zerbino DR, Birney E. Velvet: algorithms for de novo short read assembly using de Bruijn graphs. Genome research. 2008;18(5):821–9. Epub 2008/03/20. doi: 10.1101/gr.074492.107. PubMed PMID: 18349386; PubMed Central PMCID: PMC2336801.

30. Hagberg AA, Schult DA, Swart PJ, editors. Exploring Network Structure, Dynamics, and Function using NetworkX. Proceedings of the 7th Python in Science Conference; 2008; Pasadena, CA USA.

31. Pritchard L, Glover RH, Humphris S, Elphinstone JG, Toth IK. Genomics and taxonomy in diagnostics for food security: soft-rotting enterobacterial plant pathogens. Anal Methods. 2016;8(1):12–24. doi: 10.1039/c5ay02550h.

32. Gruning B, Dale R, Sjodin A, Chapman BA, Rowe J, Tomkins-Tinch CH, et al. Bioconda: sustainable and comprehensive software distribution for the life sciences. Nature methods. 2018;15(7):475–6. Epub 2018/07/04. doi: 10.1038/s41592-018-0046-7. PubMed PMID: 29967506.

33. The Anaconda Team. Anaconda Software Distribution. 2-2.4.0 ed2016.

34. van Rossum G. Python tutorial, Technical Report CS-R9526. Amsterdam: Centrum voor Wiskunde en Informatica (CWI), 1995.

35. van der Walt S, Colbert C, Varoquaux G. The NumPy Array: A structure for Efficient Numerical Computation. Computing In Science & Engineering. 2011;13:22–30.

36. Darling AE, Mau B, Perna NT. progressiveMauve: multiple genome alignment with gene gain, loss and rearrangement. PloS one. 2010;5(6):e11147. Epub 2010/07/02. doi: 10.1371/journal.pone.0011147. PubMed PMID: 20593022; PubMed Central PMCID: PMC2892488.

37. Altschul SF, Gish W, Miller W, Myers EW, Lipman DJ. Basic Local Alignment Search Tool. J Mol Biol. 1990;215(3):403–10.

38. Page AJ, Taylor J, Keane JA. Multilocus sequence typing by blast from de novo assemblies against PubMLST. The Journal of Open Source Software. 2016;1(8). doi: 10.21105/joss.00118.

39. Ondov BD, Treangen TJ, Melsted P, Mallonee AB, Bergman NH, Koren S, et al. Mash: fast genome and metagenome distance estimation using MinHash. Genome Biol. 2016;17(1):132. Epub 2016/06/22. doi: 10.1186/s13059-016-0997-x. PubMed PMID: 27323842; PubMed Central PMCID: PMCPMC4915045.

40. Hyatt D, Chen GL, Locascio PF, Land ML, Larimer FW, Hauser LJ. Prodigal: prokaryotic gene recognition and translation initiation site identification. BMC bioinformatics. 2010;11:119. Epub 2010/03/10. doi: 10.1186/1471-2105-11-119. PubMed PMID: 20211023; PubMed Central PMCID: PMC2848648.

41. Lagesen K, Hallin P, Rodland EA, Staerfeldt HH, Rognes T, Ussery DW. RNAmmer: consistent and rapid annotation of ribosomal RNA genes. Nucleic Acids Res. 2007;35(9):3100–8. Epub 2007/04/25. doi: 10.1093/nar/gkm160. PubMed PMID: 17452365; PubMed Central PMCID: PMC1888812.

42. Laslett D, Canback B. ARAGORN, a program to detect tRNA genes and tmRNA genes in nucleotide sequences. Nucleic Acids Res. 2004;32(1).

43. Bland C, Ramsey TL, Sabree F, Lowe M, Brown K, Kyrpides NC, et al. CRISPR recognition tool (CRT): a tool for automatic detection of clustered regularly interspaced palindromic repeats. BMC bioinformatics. 2007;8:209. Epub 2007/06/20. doi: 10.1186/1471-2105-8-209. PubMed PMID: 17577412; PubMed Central PMCID: PMC1924867.

44. Hunter S, Jones P, Mitchell A, Apweiler R, Attwood TK, Bateman A, et al. InterPro in 2011: new developments in the family and domain prediction database. Nucleic Acids Res. 2012;40(Database issue):D306–12. Epub 2011/11/19. doi: 10.1093/nar/gkr948. PubMed PMID: 22096229; PubMed Central PMCID: PMC3245097.

45. Claudel-Renard C, Chevalet C, Faraut T, Khan D. enzyme-specific profiles for genome annotation - PRIAM. Nucleic Acids Reseach. 2003;31(22):6633–9.

46. Yin Y, Mao X, Yang J, Chen X, Mao F, Xu Y. dbCAN: a web resource for automated carbohydrate-active enzyme annotation. Nucleic Acids Res. 2012;40(Web Server issue):W445–51. doi: 10.1093/nar/gks479. PubMed PMID: 22645317; PubMed Central PMCID: PMCPMC3394287.

47. The Gene Ontology Consortium. gene ontology: tool for the unification of biology. Nat Genet. 2000;25:95–8.

48. Mukherjee S, Huntemann M, Ivanova N, Kyrpides NC, Pati A. Large-scale contamination of microbial isolate genomes by Illumina PhiX control. Stand Genomic Sci. 2015;10:18. doi: 10.1186/1944-3277-10-18. PubMed PMID: 26203331; PubMed Central PMCID: PMCPMC4511556.

49. Sekulovic O, Fortier LC. Global transcriptional response of Clostridium difficile carrying the CD38 prophage. Appl Environ Microbiol. 2015;81(4):1364–74. doi: 10.1128/AEM.03656-14. PubMed PMID: 25501487; PubMed Central PMCID: PMCPMC4309704.

50. Horgan M, O’Sullivan O, Coffey A, Fitzgerald GF, van Sinderen D, McAuliffe O, et al. Genome analysis of the Clostridium difficile phage PhiCD6356, a temperate phage of the Siphoviridae family. Gene. 2010;462(1-2):34–43. doi: 10.1016/j.gene.2010.04.010. PubMed PMID: 20438817.

51. Bauer MP, Notermans DW, van Benthem BH, Brazier JS, Wilcox MH, Rupnik M, et al. Clostridium difficile infection in Europe: a hospital-based survey. Lancet. 2011;377(9759):63–73. doi: 10.1016/S0140-6736(10)61266-4. PubMed PMID: 21084111.

52. Knetsch CW, Terveer EM, Lauber C, Gorbalenya AE, Harmanus C, Kuijper EJ, et al. Comparative analysis of an expanded Clostridium difficile reference strain collection reveals genetic diversity and evolution through six lineages. Infect Genet Evol. 2012;12(7):1577–85. doi: 10.1016/j.meegid.2012.06.003. PubMed PMID: 22705462.

53. Dingle KE, Elliott B, Robinson E, Griffiths D, Eyre DW, Stoesser N, et al. Evolutionary history of the Clostridium difficile pathogenicity locus. Genome Biol Evol. 2014;6(1):36–52. doi: 10.1093/gbe/evt204. PubMed PMID: 24336451; PubMed Central PMCID: PMCPMC3914685.

54. Haraldsen JD, Sonenshein AL. Efficient sporulation in Clostridium difficile requires disruption of the sigmaK gene. Mol Microbiol. 2003;48(3):811–21. PubMed PMID: 12694623.

55. Liu F, Lee H, Lan R, Zhang L. Zonula occludens toxins and their prophages in Campylobacter species. Gut Pathog. 2016;8:43. doi: 10.1186/s13099-016-0125-1. PubMed PMID: 27651834; PubMed Central PMCID: PMCPMC5025632.

56. Mahendran V, Tan YS, Riordan SM, Grimm MC, Day AS, Lemberg DA, et al. The prevalence and polymorphisms of zonula occluden toxin gene in multiple Campylobacter concisus strains isolated from saliva of patients with inflammatory bowel disease and controls. PloS one. 2013;8(9):e75525. doi: 10.1371/journal.pone.0075525. PubMed PMID: 24086553; PubMed Central PMCID: PMCPMC3781098.

57. Shaw KJ, Rather PN, Hare RS, Miller GH. Molecular genetics of aminoglycoside resistance genes and familial relationships of the aminoglycoside-modifying enzymes. Microbiological Reviews. 1993;57(1):138.

58. Hackbarth CJ, Chambers HF. blaI and blaR1 regulate beta-lactamase and PBP 2a production in methicillin-resistant Staphylococcus aureus. Antimicrob Agents Chemother. 1993;37(5):1144–9. PubMed PMID: 8517704; PubMed Central PMCID: PMCPMC187918.

59. Pence MA, Haste NM, Meharena HS, Olson J, Gallo RL, Nizet V, et al. Beta-Lactamase Repressor BlaI Modulates Staphylococcus aureus Cathelicidin Antimicrobial Peptide Resistance and Virulence. PloS one. 2015;10(8):e0136605. doi: 10.1371/journal.pone.0136605. PubMed PMID: 26305782; PubMed Central PMCID: PMCPMC4549145.

60. Sandhu BK, Edwards AN, Anderson SE, Woods EC, McBride SM. Characterization of a beta-lactamase that contributes to intrinsic beta-lactam resistance in Clostridioides difficile. BioRxiv. 2019. doi: 10.1101/630020.

61. Hudson GA, Mitchell DA. RiPP antibiotics: biosynthesis and engineering potential. Current opinion in microbiology. 2018;45:61–9. Epub 2018/03/14. doi: 10.1016/j.mib.2018.02.010. PubMed PMID: 29533845; PubMed Central PMCID: PMCPMC6131089.

62. van Heel AJ, de Jong A, Song C, Viel JH, Kok J, Kuipers OP. BAGEL4: a user-friendly web server to thoroughly mine RiPPs and bacteriocins. Nucleic acids research. 2018;46(W1):W278–W81. Epub 2018/05/23. doi: 10.1093/nar/gky383. PubMed PMID: 29788290; PubMed Central PMCID: PMCPMC6030817.

63. Cascales E, Buchanan SK, Duche D, Kleanthous C, Lloubes R, Postle K, et al. Colicin biology. Microbiol Mol Biol Rev. 2007;71(1):158–229. doi: 10.1128/MMBR.00036-06. PubMed PMID: 17347522; PubMed Central PMCID: PMCPMC1847374.

64. van Belkum MJ, Martin-Visscher LA, Vederas JC. Structure and genetics of circular bacteriocins. Trends in microbiology. 2011;19(8):411–8. Epub 2011/06/15. doi: 10.1016/j.tim.2011.04.004. PubMed PMID: 21664137.

65. Briani F, Dehò G, Forti F, Ghisotti D. The Plasmid Status of Satellite Bacteriophage P4. Plasmid. 2001;45(1):1–17. doi: https://doi.org/10.1006/plas.2000.1497.

66. Brenner S, Cesareni G, Karn J. Phasmids: hybrids between Co1E1 plasmids and E. coli bacteriophage lambda. Gene. 1982;17(1):27–44. doi: https://doi.org/10.1016/0378-1119(82)90098-1.

67. Heap JT, Pennington OJ, Cartman ST, Minton NP. A modular system for Clostridium shuttle plasmids. J Microbiol Methods. 2009;78(1):79–85. doi: 10.1016/j.mimet.2009.05.004. PubMed PMID: 19445976.

68. Brouwer MS, Roberts AP, Hussain H, Williams RJ, Allan E, Mullany P. Horizontal gene transfer converts non-toxigenic Clostridium difficile strains into toxin producers. Nat Commun. 2013;4:2601. doi: 10.1038/ncomms3601. PubMed PMID: 24131955; PubMed Central PMCID: PMCPMC3826655.

69. Smillie C, Garcillan-Barcia MP, Francia MV, Rocha EP, de la Cruz F. Mobility of plasmids. Microbiol Mol Biol Rev. 2010;74(3):434–52. doi: 10.1128/MMBR.00020-10. PubMed PMID: 20805406; PubMed Central PMCID: PMCPMC2937521.

