## Supplementary figures and images for "An *in silico* survey of *Clostridioides difficile* extrachromosomal elements"

### Supplemental Figure 1 ANI high res.pdf

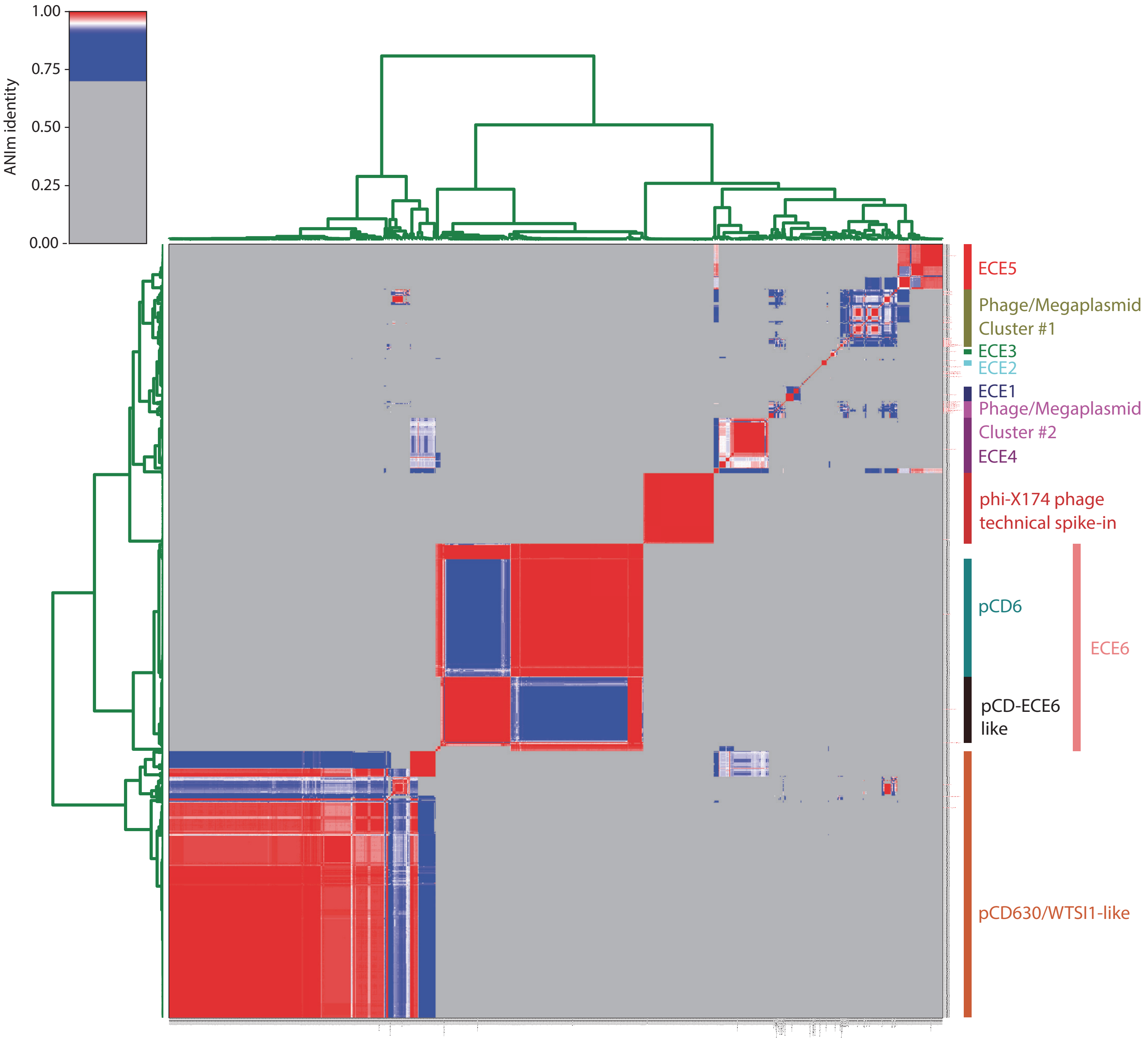
